# Tracing carbon metabolism with stable isotope metabolomics reveals the legacy of diverse carbon sources in soil

**DOI:** 10.1101/2022.04.05.487192

**Authors:** Roland C. Wilhelm, Samuel E. Barnett, Tami L. Swenson, Nicholas D. Youngblut, Chantal N. Koechli, Benjamin P. Bowen, Trent R. Northen, Daniel H. Buckley

## Abstract

Tracking the metabolic activity of whole soil communities can improve our understanding of the transformation and fate of carbon in soils. We used stable isotope metabolomics to trace ^13^C from nine labeled carbon sources into the water-soluble metabolite pool of an agricultural soil over time. Soil was amended with a mixture of all nine sources, with one source isotopically labeled in each treatment. We compared changes in the ^13^C-enrichment of metabolites with respects to carbon source and time over a 48-day incubation and contrasted differences between soluble versus insoluble sources. Whole soil metabolite profiles varied singularly by time, while the composition of ^13^C-labeled metabolites differed primarily by carbon source (R^2^ = 0.68) rather than time (R^2^ = 0.07) with source-specific differences persisting throughout incubations. The ^13^C-labeling of metabolites from insoluble carbon sources occurred at a slower rate than soluble sources but yielded a higher average atom % ^13^C in metabolite markers of biomass (amino acids and nucleic acids). The ^13^C-enrichment of metabolite markers of biomass stabilized at between 5 – 15 atom % ^13^C by the end of incubations. Temporal patterns in the ^13^C-enrichment of TCA cycle intermediates, nucleobases (uracil and thymine), and by-products of DNA salvage (allantoin) closely tracked microbial activity. Our results demonstrate that metabolite production in soils is driven by the carbon source supplied to the community, and that the fate of carbon in metabolite profiles do not tend to converge over time as a result of ongoing microbial processing and recycling.

**Importance:** Carbon metabolism in soil remains poorly described due to the inherent difficulty of obtaining information on the microbial metabolites produced by complex soil communities. Our study demonstrates the use of stable isotope probing (SIP) to study carbon metabolism in soil by tracking ^13^C from supplied carbon sources into metabolite pools and biomass. We show that differences in the metabolism of sources influences the fate of carbon in soils. Heterogeneity in ^13^C-metabolite profiles corresponded with compositional differences in the metabolically active populations, providing a basis for how microbial community composition is correlated with the quality of soil carbon. Our study demonstrates the application of SIP-metabolomics in studying soils and identifies several metabolite markers of growth, activity, and other aspects of microbial function.

## Introduction

Modeling the pathways that carbon takes through soil remains a ‘wicked’ problem because of the numerous interacting physicochemical and biological variables that regulate the metabolism of soil communities. One barrier to understanding the fate of soil carbon is our limited capacity to track the dynamic processing of carbon within soil microbial communities. The transformation of carbon via microbial metabolism impacts the residency time of carbon in soil [1], in part, due to changes in metabolite pools over time [2–5]. Yet, the mechanisms that mediate these carbon transformations remain poorly described due to the inherent difficulty of obtaining information on the microbial metabolites produced by complex soil communities.

Stable isotope probing (SIP) can be used to study the carbon metabolism of active microbial populations in soil by tracking ^13^C from supplied carbon sources into metabolite pools and biomass. Reconstructions of metabolic potential of active populations have been made from metagenome-assembled genomes with DNA-SIP [6] and from ^13^C-labeled proteins using SIP-metaproteomics [7, 8]. SIP has been used to quantify the persistence of microbial biomass carbon in soil food webs by tracking ^13^C into lipids, amino acids, and carbohydrates [9, 10]. The quantification of ^13^C in whole metabolite pools (*i*.*e*., SIP-metabolomics) has been used to study amino acid turnover in soil [11], and contrast the metabolism of glucose in oxic and anoxic soils [12], as well as in other applications [13, 14]. Here, we used SIP-metabolomics to examine community metabolism with respect to nine carbon sources that are common to soils. We did this by tracking the ^13^C-enrichment dynamics of metabolites derived from each ^13^C-labeled carbon source (‘^13^C-metabolite profiles’).

The catalytic transformation of carbon in soil is governed by the metabolic capabilities and life-history traits (*i*.*e*., growth rate, motility, morphology, etc.) represented in the soil microbial community. Life-history traits can have a greater influence over which populations metabolize common sources of soil carbon than their metabolic capacity alone (*i*.*e*., potential metabolic pathways for carbon transformation), with these traits primarily a function of growth dynamics and the bioavailability of a carbon source [15], where fast-growing bacteria dominate access to soluble sources. In addition, the biosynthetic pathways that produce cell biomass are highly conserved, and so the growth, death, and recycling of cell biomass might be expected to homogenize community metabolite pools over time regardless of carbon source [16]. Accordingly, we expected that the composition of ^13^C-metabolite pools in soils generated from ^13^C-labeled sources would vary more by time than by carbon source over the course of months.

We studied soil carbon metabolism by tracing ^13^C derived from nine isotopically-labeled carbon sources into the water-soluble soil metabolite pool across 48 days, using samples previously described in a DNA-SIP study [15]. We expected underlying differences in the metabolism of each source to be less important than the influence of growth dynamics and hypothesized that ^13^C-metabolite profiles would vary primarily with respect to time rather than carbon source. We hypothesized that any initial variation in ^13^C-metabolite profiles among carbon sources, due to differences in the composition of metabolically active bacterial populations or differences in entry point into central metabolism, would disappear over time as carbon was recycled through commonly shared pathways. We also hypothesized that ^13^C-metabolite profiles would differ between soluble (ex. glucose, xylose, amino acids, etc.) and insoluble carbon sources (cellulose and palmitic acid) due to the temporal dynamics of colonizing and degrading insoluble carbon sources [17, 18]. Finally, we examined the chemodiversity and persistence of ^13^C-enriched metabolites over time to provide an account of the *in vivo* microbial community metabolism of soil carbon.

## Materials and Methods

### Experimental design and ^13^C-labeled carbon sources

Soil was collected from an organically managed farm field in Penn Yan, New York, USA as previously described [19]. Using a microcosm approach, soil was amended with a single bolus of carbon consisting of nine carbon sources representative of soluble (amino acids, glucose, xylose, glycerol, lactate, oxalate and vanillin) and insoluble (palmitic acid and cellulose) substrates as previously described [15]. All nine carbon sources were added together and only the identity of the single ^13^C-labeled substrate was varied between treatments to allow for stable isotope probing of the metabolome over time. ^12^C-control microcosms contained all unlabelled carbon sources. Triplicate microcosms for each of the nine carbon sources were destructively sampled over time (1, 3, 6, 14, 30, and 48 days after amendment, n = 104 samples), freezing immediately at -70°C until use. Sampling times were determined by mineralization rates to capture time periods before, during, and after the period of maximal mineralization for each carbon source (overview in Table S1). Sampling was discontinued after day 14 for treatments that received soluble ^13^C-labeled sources, because these carbon sources were mineralized rapidly and the production of ^13^CO_2_ from these sources was minimal after day 14. ^12^C-control microcosms were sampled at all six timepoints.

Each carbon source was added at 0.4 mg · C g^-1^ soil for a total addition of 3.6 mg · C g^-1^ soil. Soils contained 12.2 mg · C g^-1^ dry soil of pre-existing total carbon [19]. The following ^13^C-labeled carbon sources were used: D-glucose (99 atom % ^13^C; Cambridge Isotopes, Tewksbury, MA, USA), D-xylose (99 atom % ^13^C; Omicron South Bend, IN, USA), glycerol (99 atom % ^13^C; Sigma Isotec Miamisburg, OH, USA), lactate (99 atom % ^13^C; Sigma Isotec), oxalate (99 atom % ^13^C; Sigma Isotec), palmitic acid (99 atom % ^13^C; Sigma Isotec), algal amino acid mixture (‘AA’, 98 atom % ^13^C; Cambridge Isotopes), ring-labelled vanillin (99% atom % ^13^C_6_ + 1.1% atom % ^13^C_2_; Sigma Isotec) and bacterial cellulose (∼99 atom % C) was produced in-house from the growth of *Gluconoacetobacter xylinus* on minimal media with ^13^C glucose as the sole carbon source as previously described [17]. The AA mixture derived from algae was deficient in glutamine, asparagine, cysteine, and tryptophan. Identical unlabeled carbon sources (∼1.1% atom % ^13^C) were sourced from the same company or produced in the same manner.

### Extraction of metabolites in soil water extracts

Soil metabolites were extracted following established protocols described in detail elsewhere [20]. In brief, 2 g dry weight of soil was added to 8 mL LC/MS grade water in a 15 mL conical tube and shaken on an orbital shaker for 1 hr at 200 rpm at 4°C. Samples were then centrifuged at 3220 g for 15 min at 4° C, the supernatant decanted into a 10 mL syringe attached to a 0.45 um filter disc, and subsequently filtered into 15 mL conical tubes. Filtered extracts were then lyophilized and stored at -80° C prior to LC-MS/MS analysis.

### LC-MS/MS analysis of soil metabolites

Lyophilized soil water extracts were resuspended in methanol (150 uL) containing internal standards, as previously described [21], vortexed then filtered through 0.22 um centrifugal membranes (Nanosep MF, Pall Corporation, Port Washington, NY). Buffer control samples were prepared by performing the same extraction and resuspension procedure on identical tubes without soil samples. LC/MS was performed using a ZIC-HILIC column (100 mm × 2.1 mm, 3.5 μm, 200 Å, Millipore). An Agilent 1290 series UHPLC (Agilent Technologies, Santa Clara, California, USA) was used for metabolite separation. Mobile phases were: 5 mM ammonium acetate in water (A) and 95% acetonitrile, 5% 100 mM ammonium acetate in water (B). The gradient used was: 100% B for 1.5 min (at 0.45 mL/min), linear decrease to 65% B by 15 min (at 0.45 mL/min), then decrease to 0% B by 18 min (at 0.6 mL/min), held until 23 min (at 0.6 mL/min) then returned to the initial mixture by 25 min (at 0.45 mL/min). The total runtime was 30 min. Column temperature was 40 °C. Negative and positive mode data were collected at a mass range of 70–1050 m/z in centroid data mode on a Thermo QExactive (Thermo Fisher Scientific, Waltham, MA). Fragmentation spectra (MS/MS) were acquired for some metabolites using collision energies of 10–40 eV. Sample internal standards and quality control mixtures were run at the beginning (in triplicate), end (in triplicate) and individually interspersed every 15 samples to monitor instrument performance.

### Metabolite identification

An untargeted characterization of all features evident in LC-MS/MS data was performed to estimate the overall chemical diversity of metabolite pools. Features were identified and annotated with MZmine (v. 2.23) [22]. The ‘run’ parameters have been provided in a xml file in the Supplementary Data. A targeted characterization of metabolites was performed using an in-house retention time and fragmentation library with positive assignments henceforth referred to as ‘metabolites.’ Metabolite identifications were based on comparisons with the retention times, accurate mass, and fragmentation patterns of chemical standards previously analyzed using the same column, gradient, and LC-MS/MS instrumentation. A summary of identification characteristics included similarity in fragmentation pattern, retention time, and precursor m/z, which have been provided in Table S2. LC-MS/MS data was analysed using the Metabolite Atlas [23] for the metabolite feature extraction and annotation for all targeted metabolites with an intensity ≥ 1500. Calculation of % atom ^13^C was exclusively performed on metabolites identified in the targeted analysis without interfering signals for isotopologues in the retention time integration window (see extracted ion chromatograms in Supplementary Data).

### Calculation of % atom ^13^C of metabolites

Atom % ^13^C served as a measure of the balance of source-derived carbon in metabolite pools. An increase in atom % ^13^C reflects the channeling of ^13^C into a given metabolite pool, while a decrease can indicate a pool dilution (channeling of ^12^C from unlabeled carbon sources into a given metabolite pool). Changes in atom % ^13^C may also reflect metabolite losses from the differential consumption of a ^13^C-versus ^12^C-enriched metabolites among segregated cells / populations, or vice versa. Thus, atom % ^13^C represents the relative balance between source-derived carbon entering a pool versus its dilution or loss. We did not obtain information about the absolute changes in metabolite concentrations or ^13^C-enrichment. We interpreted any lack of change in atom % ^13^C enrichment over time as an indication that this metabolite was no longer being consumed or synthesized. The % atom ^13^C of each metabolite was calculated by apportioning the peak area of each isotopologue using the following equations:

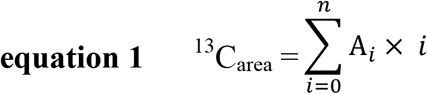

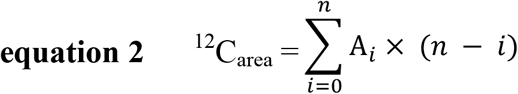

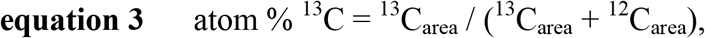

where *n* corresponds with the total number of carbon atoms in the metabolite, *i* corresponds with the number of atoms of ^13^C in each isotopologue of the metabolite, and A_*i*_ corresponds to the peak area of each isotopologue. The contribution to mass by other naturally occurring isotopologues (*e*.*g*., ^15^N, ^17^O, or ^2^H) was assumed to be negligible based on their relatively low natural abundances (0.36%, 0.038%, and 0.016%, respectively) and low occurrence in the primarily carbon-based metabolites identified. Interference in ion spectra was an issue, leading to the removal of any metabolite that had an atom % ^13^C > 0.08 in more than 2 of the 17 unlabeled ^12^C-control samples.

### Statistical analyses

All statistical analyses were performed in R (v. 3.4.0) [24]. Multiple pairwise comparisons based on Kruskal-Wallis test using ‘kruskalmc’ from the R package *pgirmess* (v. 1.7). Metabolites were deemed ^13^C-enriched based on Wilcoxon tests comparing soils amended with ^12^C-versus ^13^C-labeled carbon sources (triplicates) by day according to *p*_adj._ < 0.05. *p* values were adjusted using Benjamini-Hochberg multiple test correction (n = 612). The similarity among metabolite profiles was visualized using the t-distributed stochastic neighbor embedding (t-SNE) algorithm [25] (v. 0.15) and plotted with the R package *ggplot2* (v. 3.3.5) [26]. The ‘geom_tile’ function from *ggplot2* was used to create a heatmap of all 13C-enriched metabolites. Bacterial community composition was assessed using 16S rRNA gene amplicon count data sourced from the NCBI Short Read Archive (BioProject: PRJNA668741) and processed as described by [15]. All scripts and data required to reproduce this work are provided in the Supplementary Data can be accessed through the Open Science Foundation (doi: 10.17605/OSF.IO/V32KM).

## Results and Discussion

We profiled the community metabolism of nine ^13^C-labeled carbon sources in soil by measuring the ^13^C-enrichment of water-extractable metabolites. Carbon sources were chosen to represent a range of compound classes common in soil, and they were selected to vary widely in solubility and bioavailability [15]. A total of 2,003 features and 138 metabolites were identified using untargeted and targeted approaches, respectively. Of the total 138 metabolites, 69 yielded sufficiently interference-free m/z spectra to calculate the atom % ^13^C (Table S3). Most of the ^13^C-labeled metabolites belonged to central pathways involved in amino acid synthesis or catabolism (n = 23) and DNA synthesis (n = 10). Fifteen ^13^C-labeled metabolites corresponded to the carbon sources that were added, namely 14 amino acids and vanillin, and were enriched on day 1. A total of 67 metabolites exhibited significant ^13^C-enrichment relative to ^12^C-controls for at least a single carbon source, while 57 exhibited ^13^C-enrichment on five or more sources, and 34 were ^13^C-enriched in every metabolite profile (Table S4).

### Patterns in ^13^C-metabolites by carbon source and time

The mixture of carbon added to soil was identical in all aspects except for the identity of the ^13^C-labeled source; thus, the overall metabolite profiles should not vary based by total peak area (*i*.*e*., without consideration of isotopologues). As expected, metabolite profiles based on total peak area did not vary significantly by carbon source but did change significantly over time (Figure 1A & 1B). Temporal changes were similarly responsible for the most variation in whole bacterial community composition (Table S5) and in features in our untargeted approach (Figure S1). Notably, the total number of features significantly decreased over time (Figure 1C), suggesting a reduction in metabolite richness as metabolic products are consumed by the community. This decline in molecular diversity differs from results observed during metabolism of dissolved organic carbon in a marine system in which molecular richness increased over time [27], and this result is consistent with the expectation that carbon sources are rapidly depleted in soil, where carbon is limiting [28–30]. This observation does not necessarily fit with the hypothesis that microbial processing increases soil carbon persistence by increasing chemodiversity [5]. That said, it is likely that our method underestimates the richness of metabolites present at very low concentration.

**Figure 1.**
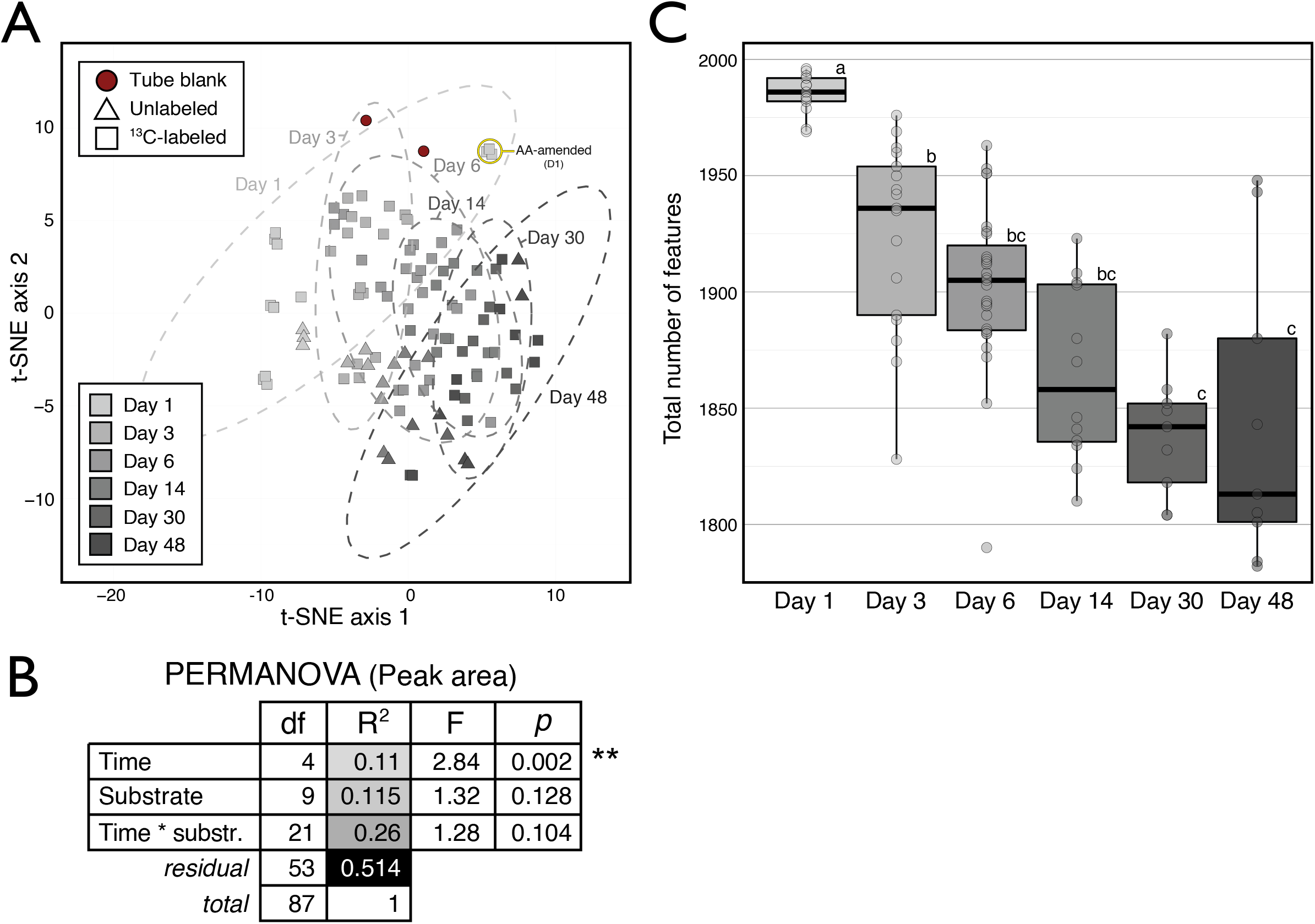
Whole metabolite profiles varied primarily across time when compared based on the peak area of all identified metabolites in soil water extracts (n = 138) irrespective of their ^13^C-enrichment. This result was expected given that all every amendment contained the whole set of carbon sources, varying solely by which single source was ^13^C-labeled. In (A), the clustering of samples based on the t-SNE multi-dimension reduction algorithm separates along the first axis according to time. The same temporal trend was apparent in untargeted features data shown in Figure S1. In (B), PERMANOVA results show incubation time was the only significant factor explaining variation in the weighted Bray-Curtis dissimilarity among metabolite profiles. Samples from day 1 were removed due to the distorting effect of ^13^C-labled amino acids (highlighted in panel A). In (C), the total number of features identified in the untargeted analysis decreased over time.

In contrast to whole feature profiles, the ^13^C-metabolite profile for each carbon source was unique and varied over time (Figure 2). The ^13^C-metabolite profiles primarily varied by carbon source (Figure 3A), which explained significantly more variation in ^13^C-enrichment than time (Figure 3B). In this way, we falsified our hypothesis that populations with similar growth strategies would share similar ^13^C-metabolite profiles. As expected, the rate at which metabolites were ^13^C-labeled differed between soluble and insoluble carbon sources (Figure 3C), but the ^13^C-metabolite profiles among soluble carbon sources did not differ broadly based on the solubility of carbon sources (Figure 3D). In fact, both soluble and insoluble carbohydrates (xylose, glucose, and cellulose) yielded the greatest number of shared ^13^C-labeled metabolites (Figure 3D), despite differences in the composition of the bacterial populations incorporating ^13^C from these sources (Figure S2). Thus, we conclude that differences in the metabolic capabilities of active populations significantly influence the composition of metabolite pools, and that differences according to carbon source persist over time.

**Figure 2.**
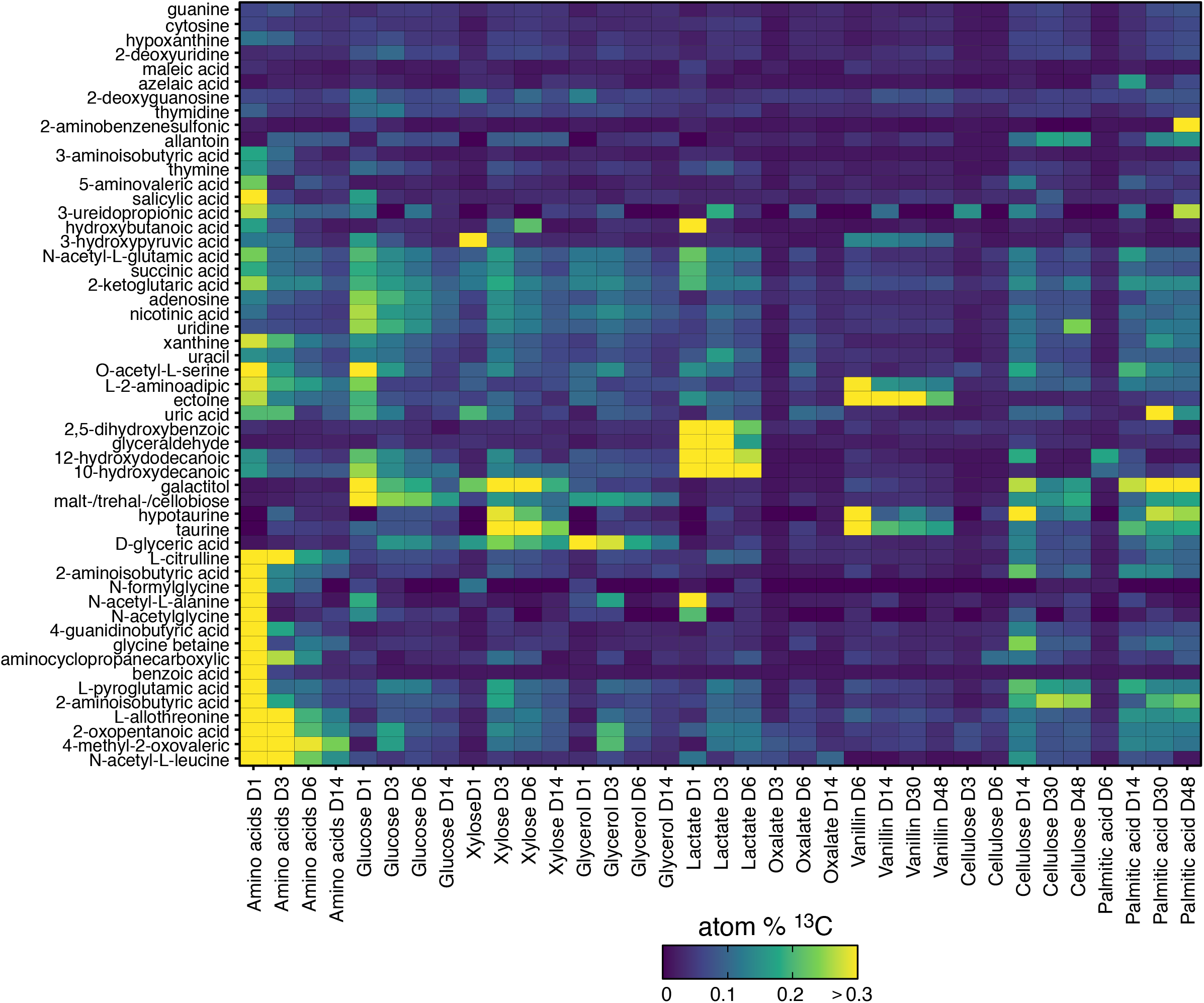
Atom % ^13^C-enrichment of metabolites varied mainly by carbon source and time. Metabolites corresponding to added ^13^C-labeled compounds were excluded. The color legend indicates degree of ^13^C-enrichment. Values represent the average of triplicates.

**Figure 3.**
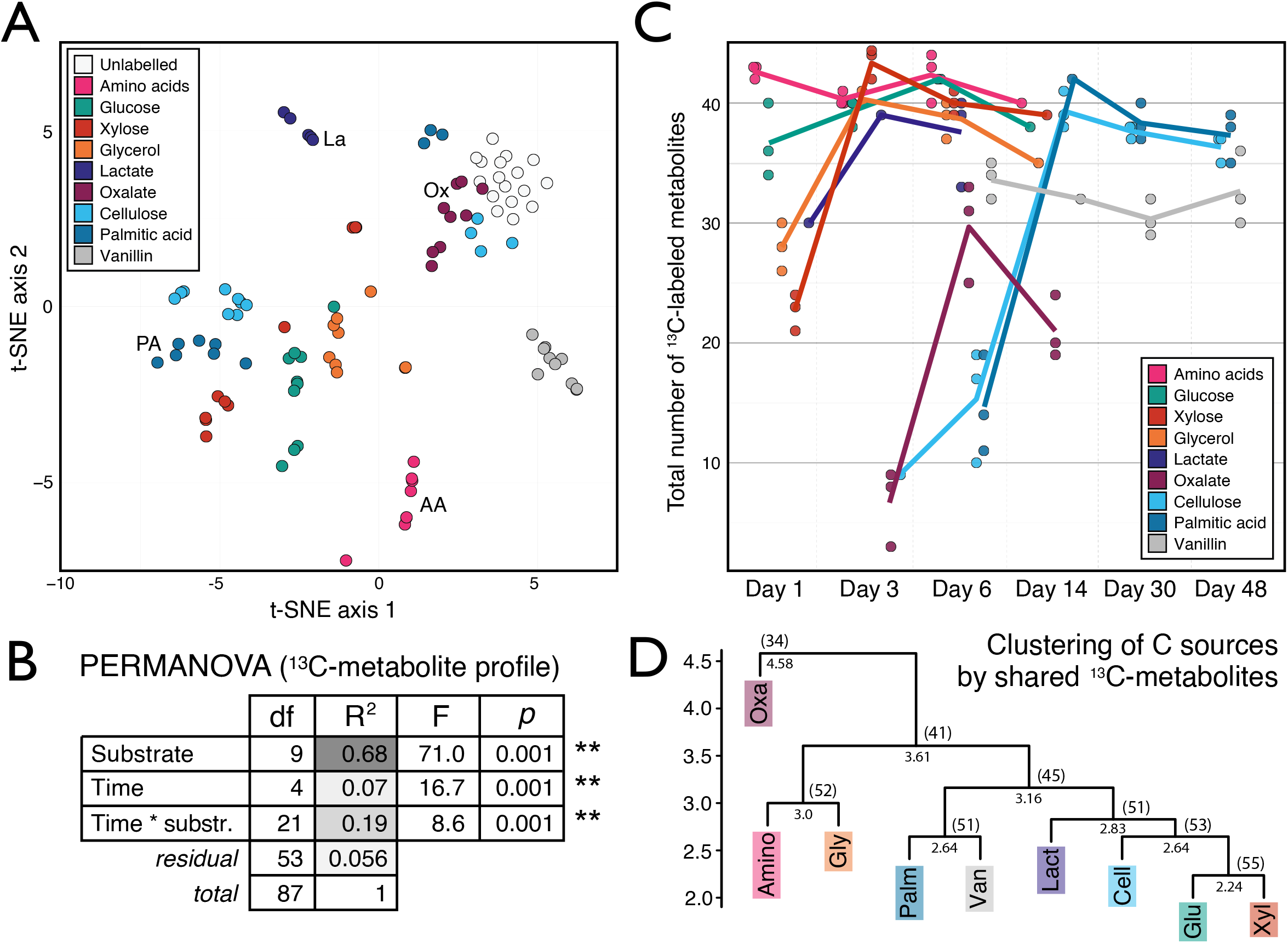
Trends in the ^13^C-enrichment of metabolites from soil water extracts varied primarily according to the ^13^C-labeled carbon source. In (A), the metabolomes clustered according to carbon source in a t-SNE based on the profile of atom % ^13^C of metabolites. To ensure clarity, some clusters are labeled with their corresponding carbon source in addition to their coloring. In (B), PERMANOVA results show carbon source explains most of the variation in the weighted Bray-Curtis dissimilarity among ^13^C-metabolite profiles. In (C), the total number of ^13^C-labeled metabolites varies over time, and these dynamics mirror ^13^C-carbon source mineralization as described for this experiment previously [15]. In (D) hierarchical clustering of metabolite profiles, based on Euclidean distance of ^13^C-metabolite presence/absence (Table S4), reveals similarities in labelling profiles between carbon sources. The cophenetic distances for each node and the number of shared metabolites were provided. Metabolites corresponding to added ^13^C-labeled compounds (n = 15) were excluded from plots except for panel (C).

### Ecological insights into soil community metabolism

The unique ^13^C-enrichment of metabolites by certain carbon sources revealed differences in the ecological characteristics of community metabolism. Aromatic compounds (benzoate and salicylate) were heavily labeled during the metabolism of the two most rapidly respired carbon sources: amino acids and glucose (Figure S3; [15]). The unique production of ^13^C-benzoic acid one day after addition of ^13^C-labeled amino acids is likely due to benzoic acid production during phenylalanine catabolism [31]. Salicylic acid, in contrast, was produced rapidly from both ^13^C-labeled amino acids and ^13^C-glucose, and this argues for a biosynthetic, rather than catabolic, origin. Bacteria such as *Pseudomonas* and *Bacillus* are known to excrete salicylic acid [32], and members of these groups were primarily ^13^C-labeled by glucose and amino acids in this experiment [15]. It is notable that both benzoic and salicylic acids serve as phytohormones and have antifungal properties [33, 34], though the mechanism and consequence of production remains to be determined.

The ^13^C-enrichment of metabolites involved in osmoregulation (*i*.*e*., compatible solutes) differed broadly among soluble and insoluble carbon sources, and vanillin. The highest ^13^C-enrichment of glycine betaine occurred during periods of peak respiration for both soluble (day 1) and insoluble sources (day 14), while vanillin produced low levels of ^13^C-enrichment (Figure 4). In comparison, maximal ^13^C-enrichment of ectoine was observed from vanillin, with moderate and minimal enrichment resulting from soluble and insoluble sources, respectively. Ectoine can protect cells from damage caused by reactive oxygen species (ROS), whereas glycine betaine does not [35, 36]. It is possible that differences in ^13^C-labeling of ectoine may reflect the needs of cells to protect from ROS generated during the oxidative catabolism of aromatics [37–40], like vanillin, and during periods of high metabolic activity [41].

**Figure 4.**
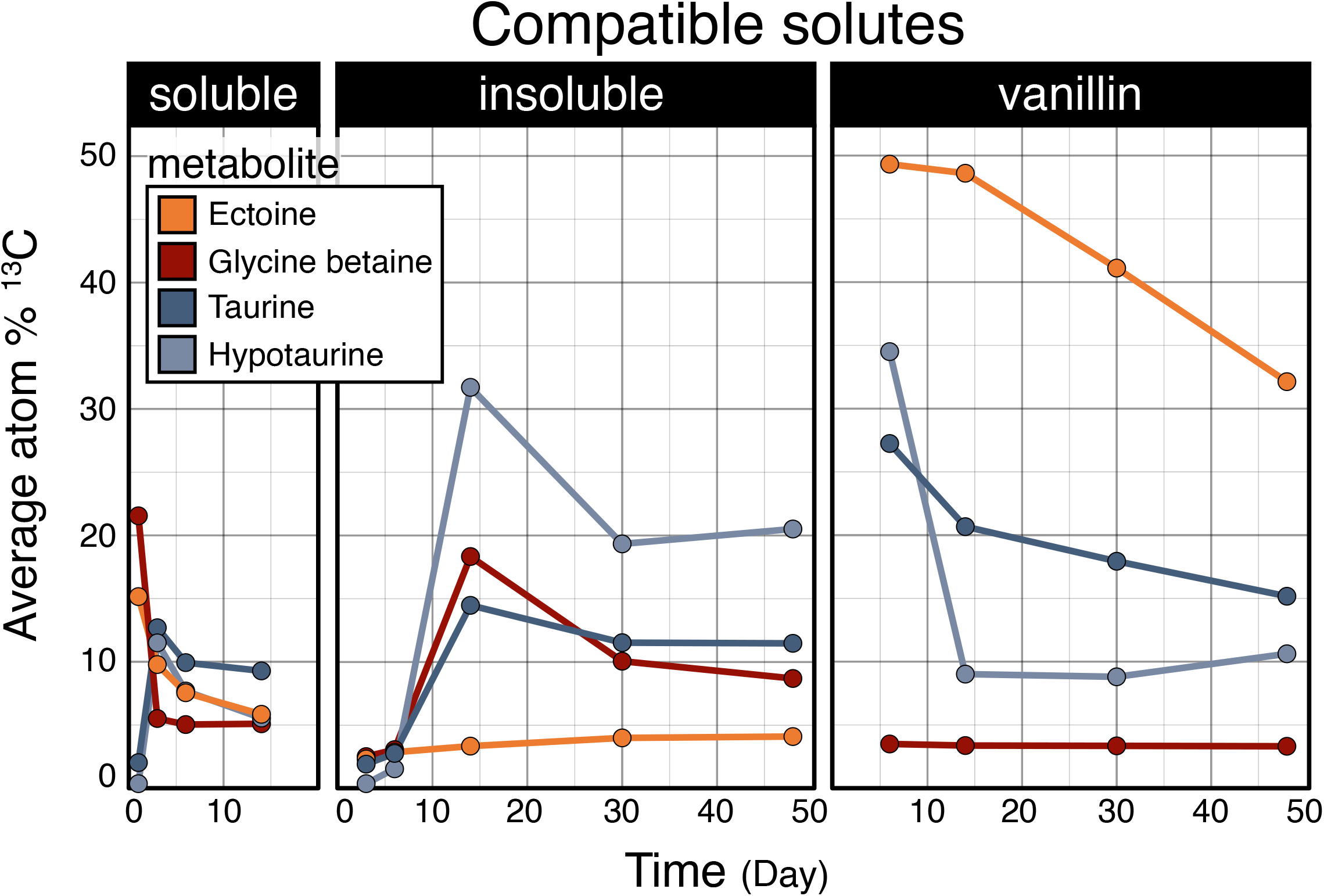
Patterns in the atom % ^13^C enrichment of metabolites identified as compatible solutes differed with respect to carbon source. Trends in major compatible solutes differ among compound types, with ectoine highly enriched during the metabolism of vanillin. Carbon sources were aggregated by response into three classes: soluble (glucose, xylose, amino acids, lactic acid, glycerol, oxalic acid), insoluble (cellulose and palmitic acid), and vanillin. Points represent average atom % ^13^C.

The ^13^C-enrichment of disaccharides (including maltose, trehalose and cellobiose) also differed between soluble and insoluble carbon sources. Disaccharide enrichment increased over time for insoluble carbon sources, while the opposite trend was generally evident for soluble sources (Figure 5). For cellulose, this trend may track the liberation of cellobiose during cellulose degradation, but this cannot explain the same trend observed for palmitic acid. Maltose and trehalose are common metabolites that serve in carbon and energy storage, enhancing survival [42, 43] and fueling growth following periods of starvation [44–46]. Our observations suggest that populations metabolizing insoluble carbon sources might favor sustained carbon and energy storage during growth, while populations that metabolize soluble carbon sources begin depleting reserves immediately following peak activity.

**Figure 5.**
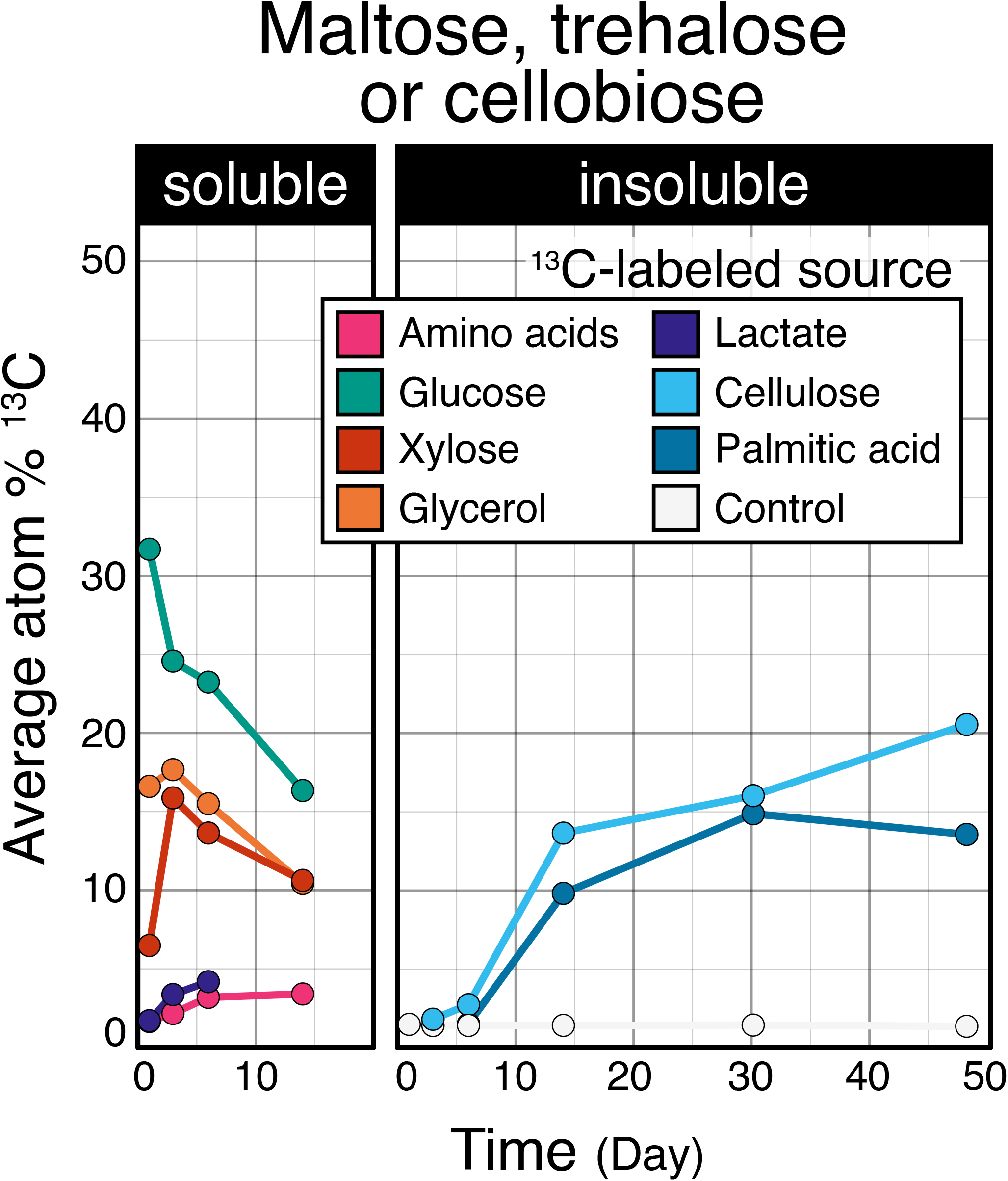
Patterns in the atom % ^13^C enrichment of metabolites identified as disaccharides differed with respect to carbon source. The proportion of source-derived ^13^C occurring in disaccharides (maltose, trehalose or cellobiose) steadily increased during the degradation of insoluble compounds. Maltose, trehalose, and cellobiose overlap in m/z and could not be distinguished using our chromatography conditions. Values represent the average of triplicates. Carbon sources were aggregated by response into two classes: soluble (glucose, xylose, amino acids, lactic acid, glycerol, oxalic acid), insoluble (cellulose and palmitic acid).

### Microbial growth and the persistence of biomass in soil

The time of peak metabolite enrichment (*i*.*e*., when the most features were ^13^C-enriched; Figure 3C) was consistent with the dynamics of ^13^CO_2_ respiration. Glucose and amino acids were mineralized earliest (< 24 hrs), followed by lactate, xylose, and glycerol (< 2 days), then oxalate (day 6), cellulose (day 10), and palmitic acid (day 14) [15]. Rapid ^13^C-enrichment and subsequent depletion of TCA cycle intermediates (*e*.*g*., succinic and 2-ketoglutaric acids), corresponded with peak mineralization activity (Figure 6A). Notably, the atom % ^13^C of 2-ketoglutaric acid was universally higher than succinic acid. Since 2-ketoglutaric acid is upstream of succinic acid and is a branch point for amino acid biosynthesis, these results likely illustrate ^13^C pool dilution as carbon passes through the TCA cycle, though isotopic fractionation might also explain differences. Similarly, the ^13^C enrichment of uracil (RNA) versus thymine (DNA) diverged most during periods of highest respiration (Figure 6B), occurring on day 3 (for all soluble carbon sources) and day 14 (insoluble sources)

**Figure 6.**
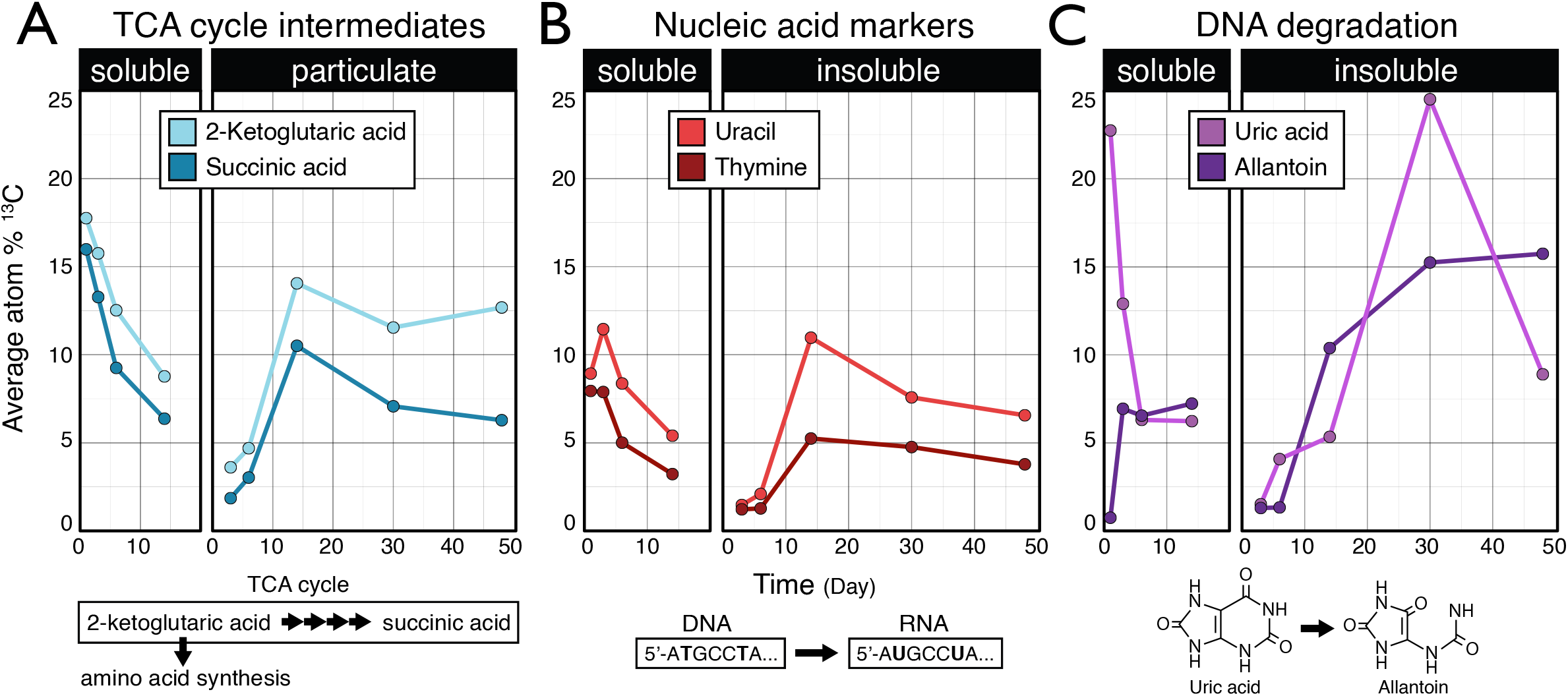
Soluble and insoluble carbon sources exhibited dramatic differences with respect to the enrichment of TCA cycle intermediates, nucleotides, and DNA degradation products. In (A), the relative ^13^C enrichment of the upstream TCA intermediate 2-ketogultaric acid (also 2-oxoglutaric acid) was universally higher than succinic acid, indicative of the shunting of carbon from 2-ketogultaric acid into amino acid synthesis. In (B), the enrichment of uracil and thymine provides data on the proportion of RNA and DNA over time. In (C), the degradation of purine bases generates uric acid, which is then cleaved to produce allantoin as part of a salvage pathway yielding glycine, urea, ammonia, and energy. Carbon sources were aggregated by response into two classes: soluble (glucose, xylose, amino acids, lactic acid, glycerol, oxalic acid), insoluble (cellulose and palmitic acid). Points represent average atom % ^13^C.

Most markers of microbial biomass and growth, such as amino acids, nucleotides, and organic acids, exhibited broadly similar patterns of ^13^C-enrichment regardless of carbon source, differing primarily with respect to carbon source solubility (Figure 7). The insoluble carbon sources, cellulose and palmitic acid, are catabolized through very different pathways, and so the similarity in ^13^C-metabolite profiles is likely due to common biosynthetic pathways shared among bacteria that colonize and degrade insoluble compounds [17, 18]. The enrichment of biomass markers attenuated over time, stabilizing at between 5 – 15 atom % ^13^C by the end of incubations (Figure 7). Amino acids had the highest level of enrichment among biomass markers at the end of every incubation (Table 1). This observation is consistent with the long residency time of amino acids in soil [47–49], and likely reflects the preservation of protein in dormant cells or in soil necromass. The latter case is supported by the disproportionately high stabilization of protein relative to other necromass fractions in soils [49]. The accumulation of allantoin, the only metabolite to consistently increase in atom % ^13^C from all carbon sources over time (Figure 6C), is further evidence of the persistence of source-derived ^13^C in necromass. Allantoin is produced as part of a salvage pathway during DNA degradation, first yielding uric acid then allantoin. Allantoin is a valuable source of energy and nitrogen to soil communities [50–52], thus its accumulation likely reflects the accumulation of ^13^C in soil necromass.

**Table 1.** Higher proportions of source-derived carbon remained in amino acids and their derivatives over time relative to organic acid and nucleic acid metabolite pools according to the average atom % ^13^C at the end of each incubation. The standard deviation (±) among triplicates is shown, except for amino acids, glucose, glycerol, and xylose because only a single sample was available at the final timepoint.

| **Carbon source** | **day** | **amino acids** | **amino acid derivatives** | **DNA** | **organic acids** |
| --- | --- | --- | --- | --- | --- |
| No label | 48 | 2.4 ± 0.04 | 1.8 ± 0.5 | 2.2 ± 0.1 | 1.6 ± 0.2 |
| Amino acids | 14 | 14.2 | 8.7 | 4.7 | 5.0 |
| Glucose | 14 | 6.6 | 4.9 | 6.2 | 4.6 |
| Glycerol | 14 | 6.3 | 4.1 | 4.6 | 4.1 |
| Xylose | 14 | 8.6 | 5.9 | 6.0 | 5.2 |
| Oxalate | 14 | 5.5 ± 1.2 | 2.8 ± 0.4 | 3.7 ± 0.3 | 2.0 ± 0.2 |
| Lactate | 6 | 8.5 ± 0.7 | 7.2 ± 0.4 | 5.6 ± 0.1 | 5.4 ± 0.5 |
| Cellulose | 48 | 11.0 ± 2.1 | 8.2 ± 1.6 | 6.8 ± 0.6 | 6.5 ± 0.07 |
| Palmitic acid | 48 | 13.1 ± 3.3 | 7.6 ± 1.8 | 6.9 ± 1.2 | 6.7 ± 2.1 |
| Vanillin | 48 | 6.6 ± 1.5 | 5.6 ± 2.5 | 4.2 ± 0.3 | 3.5 ± 0.2 |
| average atom % <sup>13</sup>C |  | 8.1 | 5.6 | 4.9 | 4.3 |

**Figure 7.**
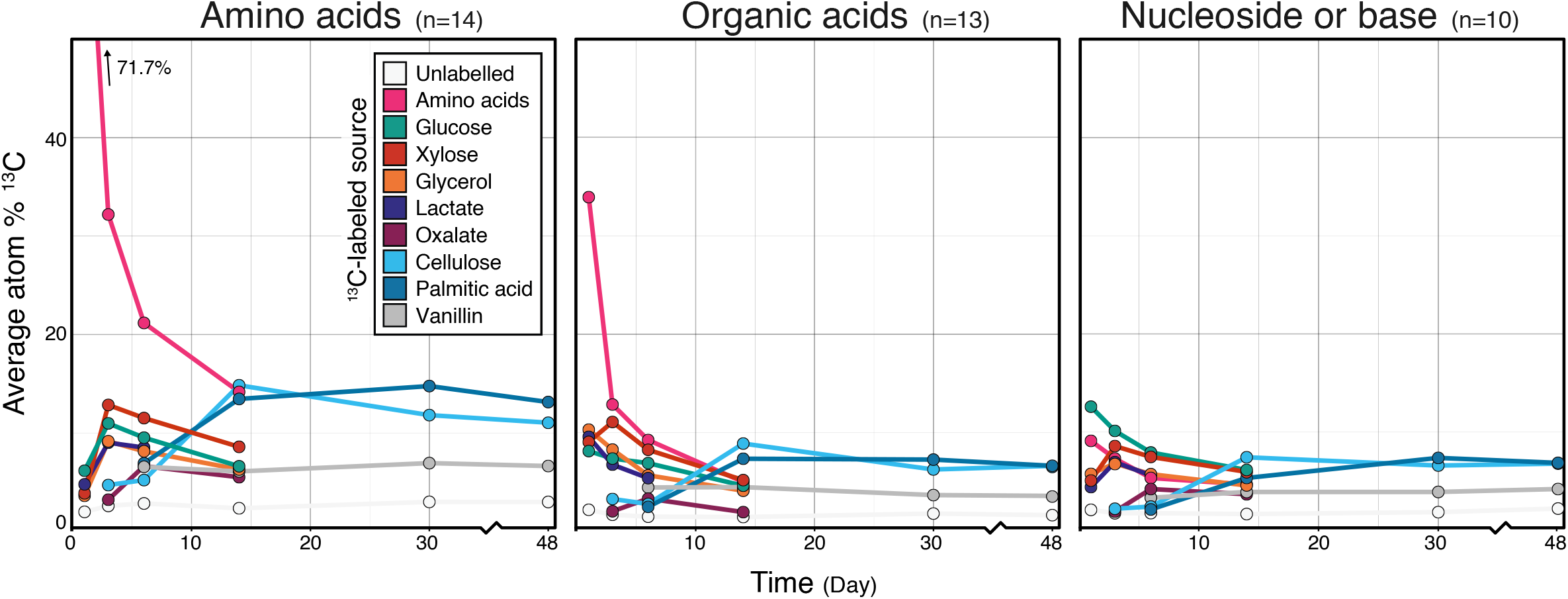
The average atom % ^13^C enrichment of amino acids, organic acids, and nucleosides/bases. The metabolites grouped into these classes are provided in Table S3. Metabolites corresponding to added amino acids were included in the average. The apparent low ^13^C enrichment of amino acids by day 1 likely reflects our co-addition of unlabeled amino acids which increased overall pool size, thereby artificially reducing the proportion of newly synthesized, ^13^C-labeled amino acids. Values represent the average of triplicates.

## Conclusions

Our study demonstrated the extent to which the metabolism of diverse carbon sources affects the fate of carbon in soils. Contrary to our expectations, ^13^C-metabolite profiles varied more by carbon source than by time, with source-specific differences persisting throughout incubations. This suggests that homogenizing forces, like biomass recycling, are less transformative of soil metabolite pools over the short-term (∼ month) than the initial community metabolism of a carbon source. Heterogeneity in ^13^C-metabolite profiles corresponded with compositional differences in the metabolically active populations [15], which provides a basis for how microbial community composition is correlated with the quality of soil carbon [53].

Our study is among the first examples of SIP-metabolomics in soil systems and demonstrates the capacity to profile metabolite markers of growth, activity, and other aspects of microbial function, such as osmoregulation. Our study was limited in the number of metabolites for which we could reliably calculate atom % ^13^C. However, tools are becoming available to assist in identifying metabolites in complex biological systems using SIP-metabolomics [54]. The full realization of SIP-metabolomics for modeling the pathways of carbon through soil may lie in a comprehensive multi-omic approach. SIP-metabolomic data can be paired with highly-resolved genomic, metaproteomic, and relative activity for microbial populations [55–59] to provide a more complete model of the soil community carbon metabolism.

## Acknowledgements

This work was supported by the U.S. Department of Energy, Office of Biological & Environmental Research Genomic Science Program under award number DE-SC0016364. Any opinions, findings, conclusions, or recommendations expressed in this publication are those of the author(s) and do not necessarily reflect the views of the United States Department of Energy.

## Author Contributions

RCW performed research and writing. RCW and SEB performed data analysis. TW performed LCMS analyses, and BPB pre-processed LCMS data. DHB and NDY designed the microcosm experiment. DHB and TRN guided all research efforts.

