## Supplementary Figures for "Tracing carbon metabolism with stable isotope metabolomics reveals the legacy of diverse carbon sources in soil"

**Figure S1**. The feature profiles of all soil water extracts varied primarily according due to incubation length when compared based on the peak area (n = 2,003). In (A), the clustering of samples based on the t-SNE multi-dimension reduction algorithm separates along the first axis according to time. In (B), PERMANOVA results show incubation time was the primary factor explaining variation in the weighted Bray-Curtis dissimilarity among metabolite profiles. In (B), Samples from day 1 were removed due to the distorting effect of $^{13}C$-labled amino acids (highlighted in panel A).

A

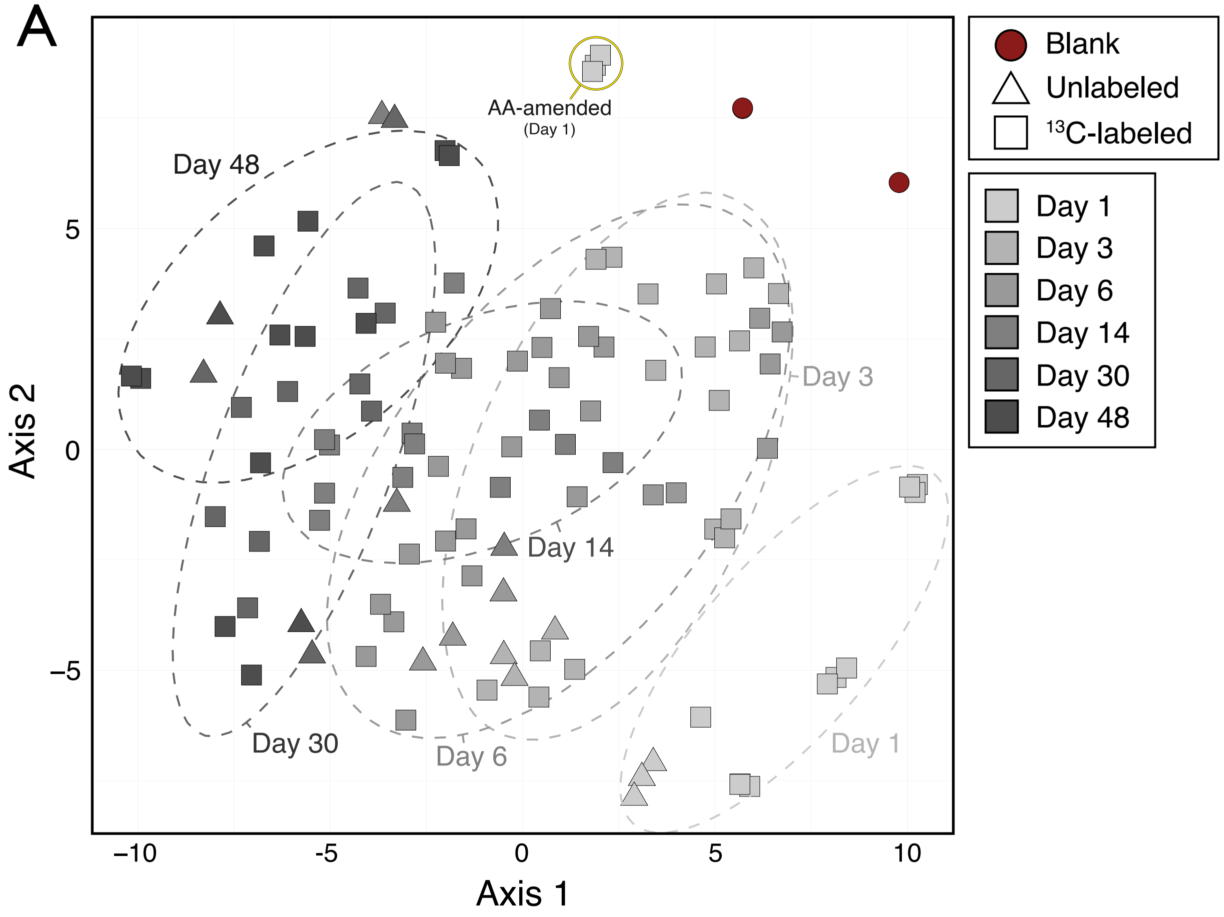


B

PERMANOVA (Peak area)

| | df | $R^2$ | F | *p* | |
|---|---|---|---|---|---|
| Time | 4 | 0.401 | 15.4 | 0.001 | *** |
| Substrate | 9 | 0.095 | 1.64 | 0.02 | * |
| Time * substr. | 21 | 0.139 | 1.02 | 0.47 | |
| *residual* | 56 | 0.364 | | | |
| *total* | 90 | 1 | | | |

**Figure S2**. An NMDS ordination showing the dissimilarity of bacterial communities which were $^{13}C$-labeled by soluble versus insoluble substrates. The differences in composition between populations metabolising glucose and cellulose are noteworthy, given the similarity in $^{13}C$-enriched metabolites shared (see Figure 2C). Bacterial community composition was determined from amplicon libraries targeting the 16S rRNA gene in the 'heavy' fractions of a DNA density gradient as described by Barnett *et al*., 2021, from which the data was obtained.

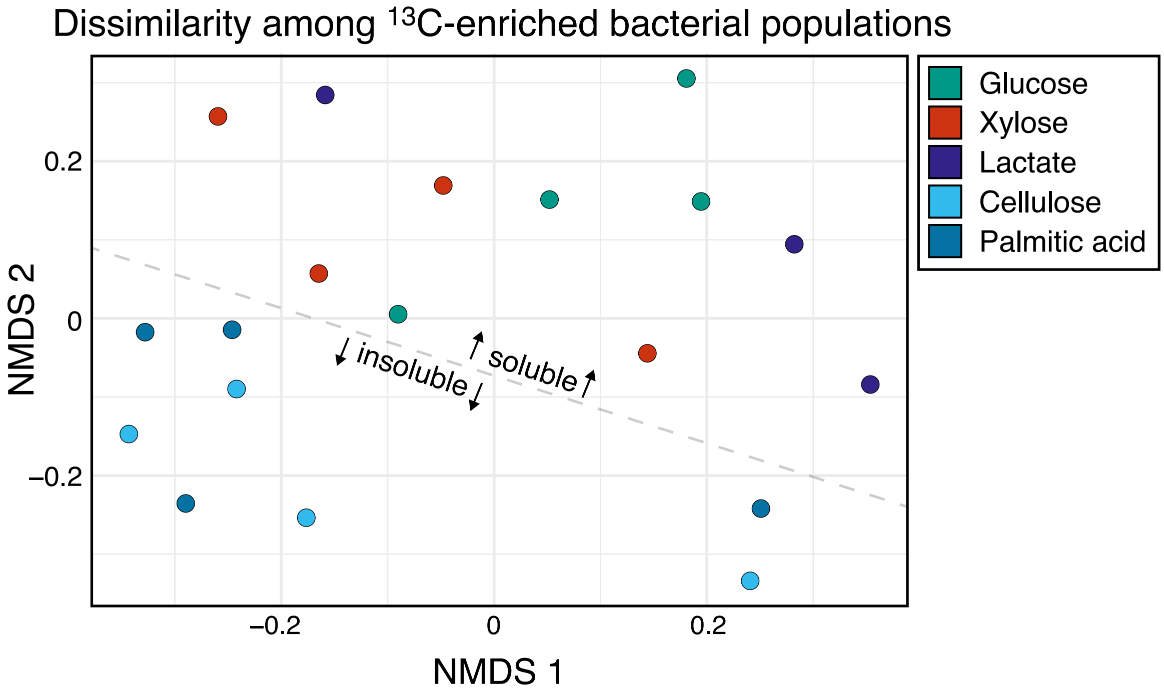

**Figure S3**. Patterns in the atom % $^{13}C$ enrichment of benoic and salicylic acid which were heavily enriched at the earliest timepoints in soils amended with amino acids and glucose.

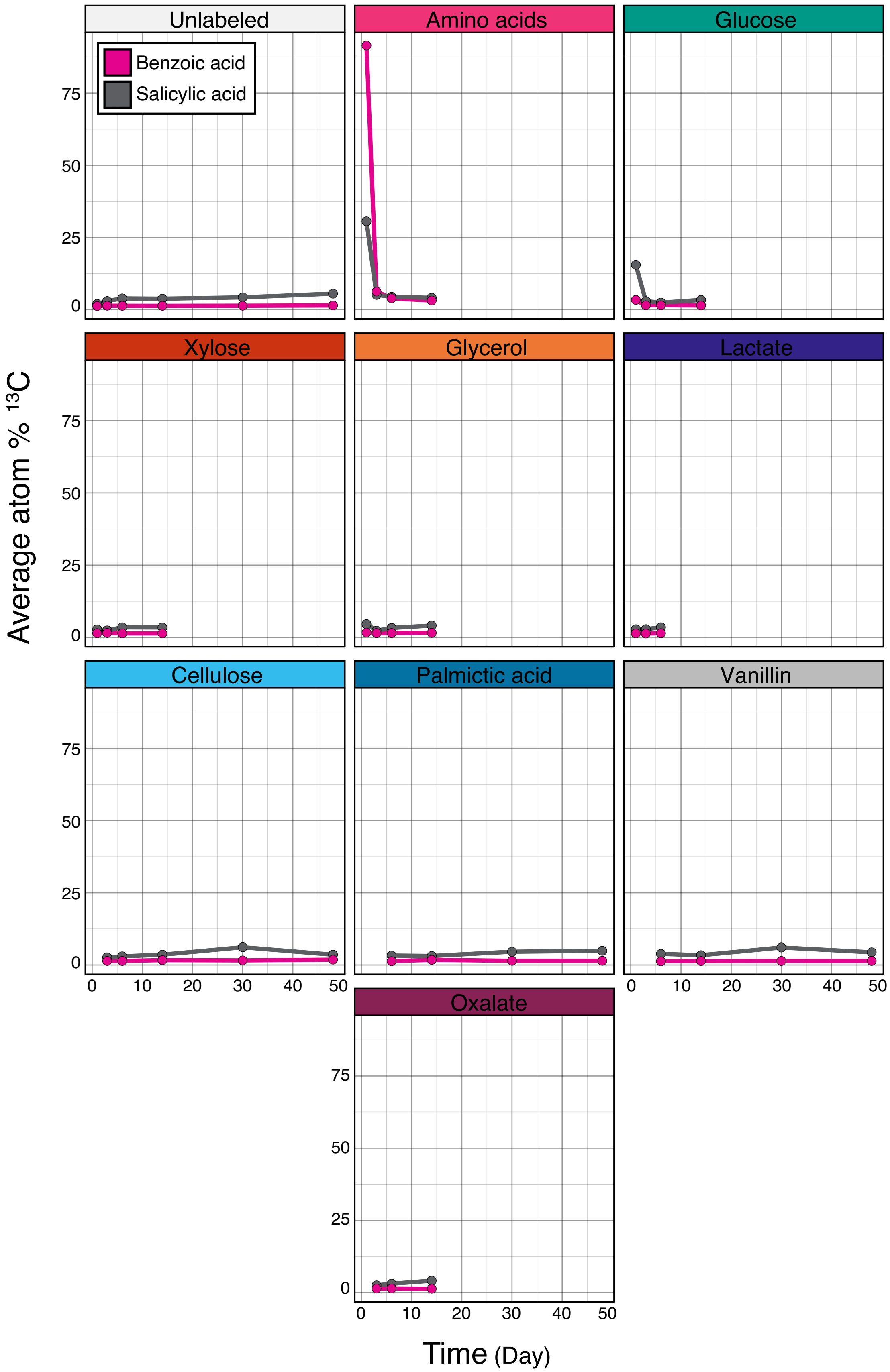
